# Predicting the evolution of lung squamous cell carcinoma in situ using deep learning

**DOI:** 10.1101/2022.12.06.519339

**Authors:** Alon Vigdorovits, Gheorghe-Emilian Olteanu, Ovidiu Tica, Monica Boros, Ovidiu Pop

## Abstract

Lung squamous cell carcinoma *in situ* (SCIS) is the pre-invasive precursor lesion of lung squamous cell carcinoma (SCC). Only half of these lesions progress to invasive cancer, while a third undergo spontaneous regression. The ability to predict the evolution of SCIS lesions can significantly impact the management of lung cancer patients.

Here, we present the use of the deep learning (DL) approach in order to predict the progression of SCIS. The dataset consisted of 112 H&E stained whole slide images (WSI) that were obtained from the Image Data Resource public repository. The data set corresponded to tumors of patients who underwent biopsies of SCIS lesions and were subsequently followed up by bronchoscopy and CT to monitor for progression to SCC. We show that a deep convolutional neural network (DCNN) can predict if a SCIS lesion will progress to SCC. The model achieved a per-tile AUC of 0.78 (SD = 0.01) on the test set, an F1 score of 0.84 (SD = 0.05), and a sensitivity of 0.94 (SD = 0.01). Class activation maps were created in order to explore how the DCNN made decisions.

To our knowledge, this study is the first to demonstrate that DL has the ability to predict the evolution of SCIS from H&E WSI. DL has the potential to be used as a low-cost method that could provide prognostic information for patients with preinvasive lesions.

## Introduction

Lung cancer is the leading cause of cancer mortality, with an estimated 1.8 million deaths per year (1). Squamous cell carcinoma of the lung (SCC) accounts for about 20% of all lung cancers and is more heavily associated with smoking than lung adenocarcinoma (2). It usually occurs in the proximal part of the airway and originates typically from the basal cells of the bronchial mucosa (3). SCC shows numerous genetic alterations but only a few clinically actionable driver mutations, in contrast to lung adenocarcinoma, which has several targetable driver mutations such as *EGFR, ALK,* and *ROS1* (4). Squamous cell carcinoma in situ (SCIS) is the preinvasive precursor lesion of SCC. Given the fact that 30% of SCIS undergo spontaneous regression, the clinical management of patients that present with SCIS is challenging and often results in overtreatment (5). Usually, patients have multiple comorbidities, further complicating clinical decisions (6).

In recent years, low-dose helical CT screening seems to offer a promising way to improve survival in SCC (7). Unfortunately, a CT scan might not always detect preinvasive lesions (5). The ability to predict which SCIS will progress to SCC would be invaluable to help guide further monitoring and treatment. Previous studies have characterized the molecular profiles of SCIS in order to predict if they will progress to SCC or spontaneously regress (8). Nonetheless, given the fact that progressive and regressive lesions are indistinguishable from each other, morphology was not directly used to predict progression.

DL is a subfield of machine learning that uses artificial neural networks (ANN) in order to learn patterns from highly complex data. ANN are non-linear statistical models that are loosely based on biological neural networks and have achieved tremendous success in various pattern recognition tasks (9). Medical imaging and especially histopathology are ideal for analysis via DL techniques due to their high information density (10). Researchers have used DL to classify various tumors, predict molecular alterations directly from H&E stained images, and estimate survival from histomorphology (11) (12) (13). Histopathology images can thus be data-mined for a wealth of clinically actionable data, some of which could hold information regarding the natural history of certain lesions.

In this study, we apply a DL approach to H&E stained images in order to predict which SCIS lesions will evolve and progress to SCC and which lesions will regress. To our knowledge, this is the first study that uses DL on pathology images in an attempt to predict the course of a preinvasive lesion. It is our hope that this study will drive other researchers to explore the use of DL on WSI in order to predict the evolution of other preinvasive precursor lesions.

## Materials and Methods

### Dataset

We used a publicly available WSI dataset from patients with SCIS (14), obtained from the Image Data Resource public repository (15). This represents the largest cohort of patients with these types of lesions. The cohort consisted of patients with SCIS that underwent conservative management, with autofluorescence bronchoscopy being performed every 4 months and CT scans every 12 months. Treatment was only performed when a SCIS lesion progressed to invasive cancer, as demonstrated by histopathology (8) (16). The dataset consisted of 112 H&E stained WSI in SVS format and the corresponding regression or progression label. Of these, 68 corresponded to lesions that progressed to invasive cancer and 44 to lesions that underwent regression.

### Image preparation

In order to build the image dataset used in training the deep convolutional neural network (DCNN), regions of interest containing the lesions were selected by a thoracic pathologist. At this point, in order to prevent data leakage generated by having images from the same patient used for both training and testing, a test dataset consisting of 20% of the WSI was generated by random sampling. Tiles of 256 x 256 pixels at 20x magnification were cropped from the WSI using a custom script (Figure 1). The number of tiles per WSI varied depending on the size of the biopsy and the size of the lesion. The final dataset contained 11130 images. Of these, 7907 images were from lesions that progressed to invasive cancer and 3223 from lesions that regressed. The training dataset had 8800 images, while the test dataset contained 2330 images. In order to improve the generalizability of the model, various data augmentation techniques such as random vertical flipping, random horizontal flipping, random rotations and color jitter were applied to the images.

**Figure 1:**
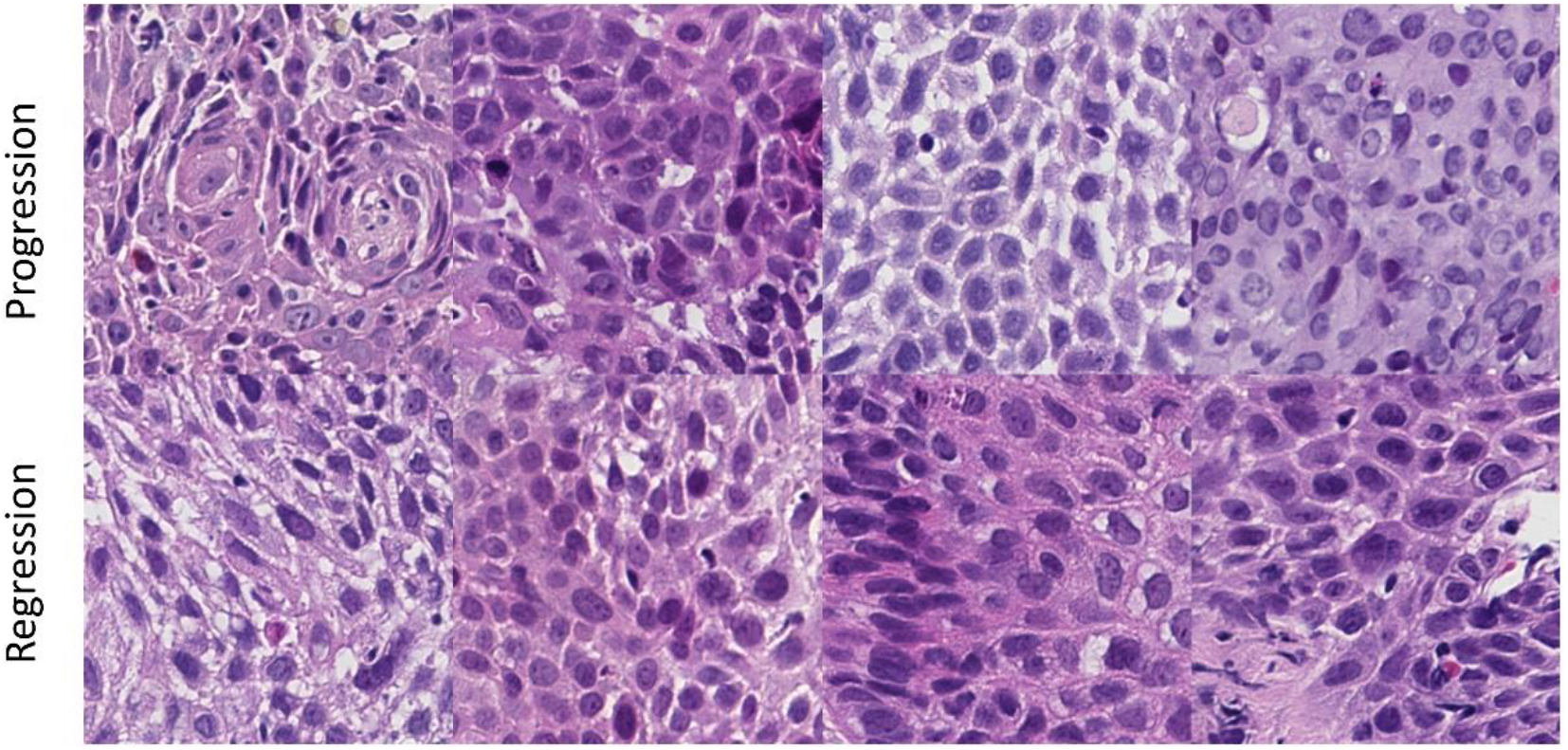
Examples of images used to train the model

### Deep learning method

The training dataset was used to train a modified ResNet18 network architecture (17). Our DL library of choice was PyTorch (18). A training/validation split of 80/20 was used, with model hyperparameters being tuned according to results obtained on the validation set. The fully connected portion of the ResNet18 model contained a dropout layer with a dropout frequency rate of 0.7 in order to prevent overfitting. The model was pretrained on ImageNet and then fine-tuned on the training data using a stochastic gradient descent optimization algorithm with a learning rate of 10^-3^ for 5 epochs, after which the learning rate was decreased to 10^-4^ for another 5 epochs (19). At this point, the model had already reached convergence. The per-tile performance metrics used were: F1 score, AUC (area under the receiver operating characteristic curve), recall (sensitivity) and precision. F1 score was included due to the limited interpretability of AUC when faced with an imbalanced classification problem. The F1 score takes into account both recall and precision and is equal to 2 x (precision x recall) / (precision + recall). The mean and standard deviation obtained across 5 random seeds were reported. We applied t-distributed Stochastic Neighbor Embedding (t-SNE) on test data to map the high dimensional information used by the DCNN onto the two-dimensional plane, in order to see how well the model separates the data. In order to gain insight into the regions in the images that are the most relevant for model decision making, we also created class activation maps (CAM) of the predicted classes (20).

### Code availability

All of the code used for image preparation as well as model training and testing is available at https://github.com/ohalon/SCISEvo.

## Results

The model achieved a mean per-tile AUC of 0.784 with a standard deviation (SD) of 0.01 (Figure 2). The selected threshold probability for progression was 0.5. The DCNN model also yielded an F1 score of 0.84 (SD = 0.05), a sensitivity (recall) of 0.94 (SD = 0.01) and a precision of 0.76 (SD = 0.007). Figure 3 represents the confusion matrix obtained from the classification results, showing a low percentage of false negatives for detecting progression, which leads to high sensitivity.

**Figure 2:**
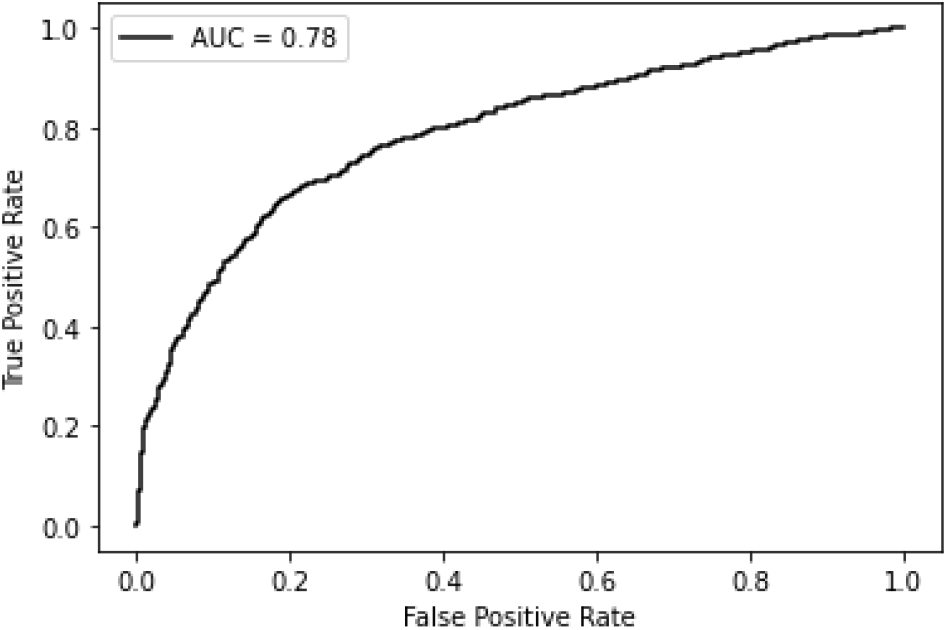
Receiver operating characteristic curve for the model

**Figure 3:**
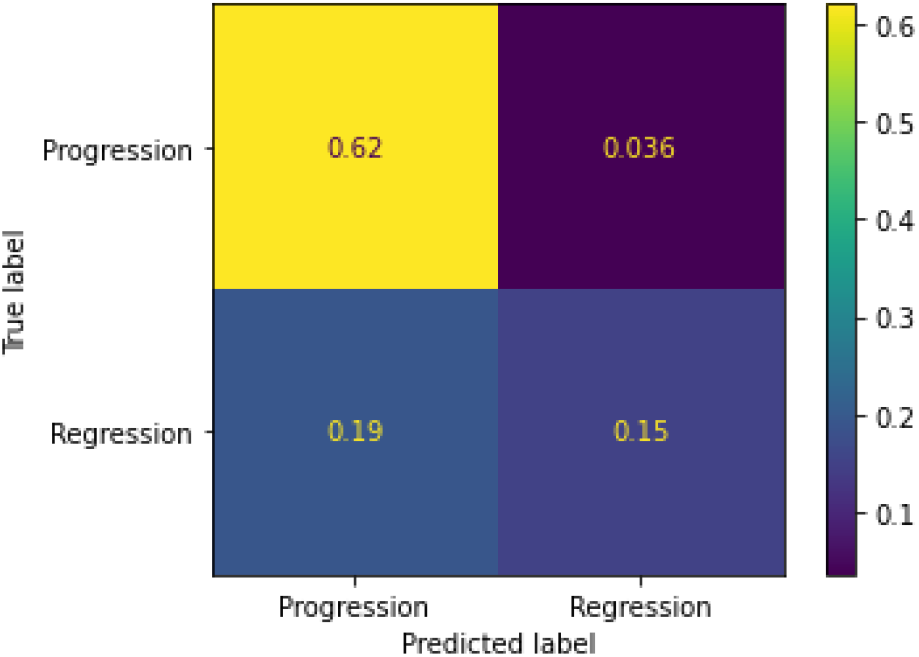
Confusion matrix of the model

The t-SNE plot revealed 2 clusters of images corresponding roughly to the progression and regression classes. The progression cluster was tighter and contained relatively few images with regression, in agreement with the model’s good performance in detecting progression. The regression cluster was more spread out and contained a large number of cases with progression, indicating the difficulty of the model in separating the regression images (Figure 4).

**Figure 4:**
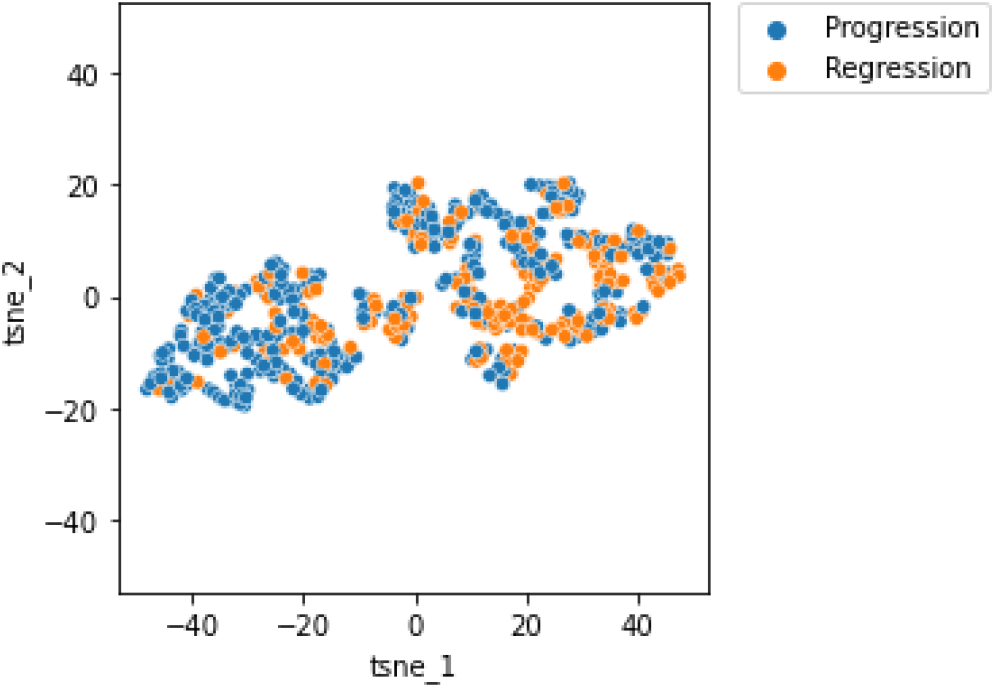
t-SNE plot for the model classification

CAMs were created for a number of random examples of both correct and incorrect classifications. These activation maps can aid in obtaining discriminative image regions that are used by the DCNN to classify an image. Areas colored in yellow represent regions of the image that are deemed important by the model (Figure 5).

**Figure 5:**
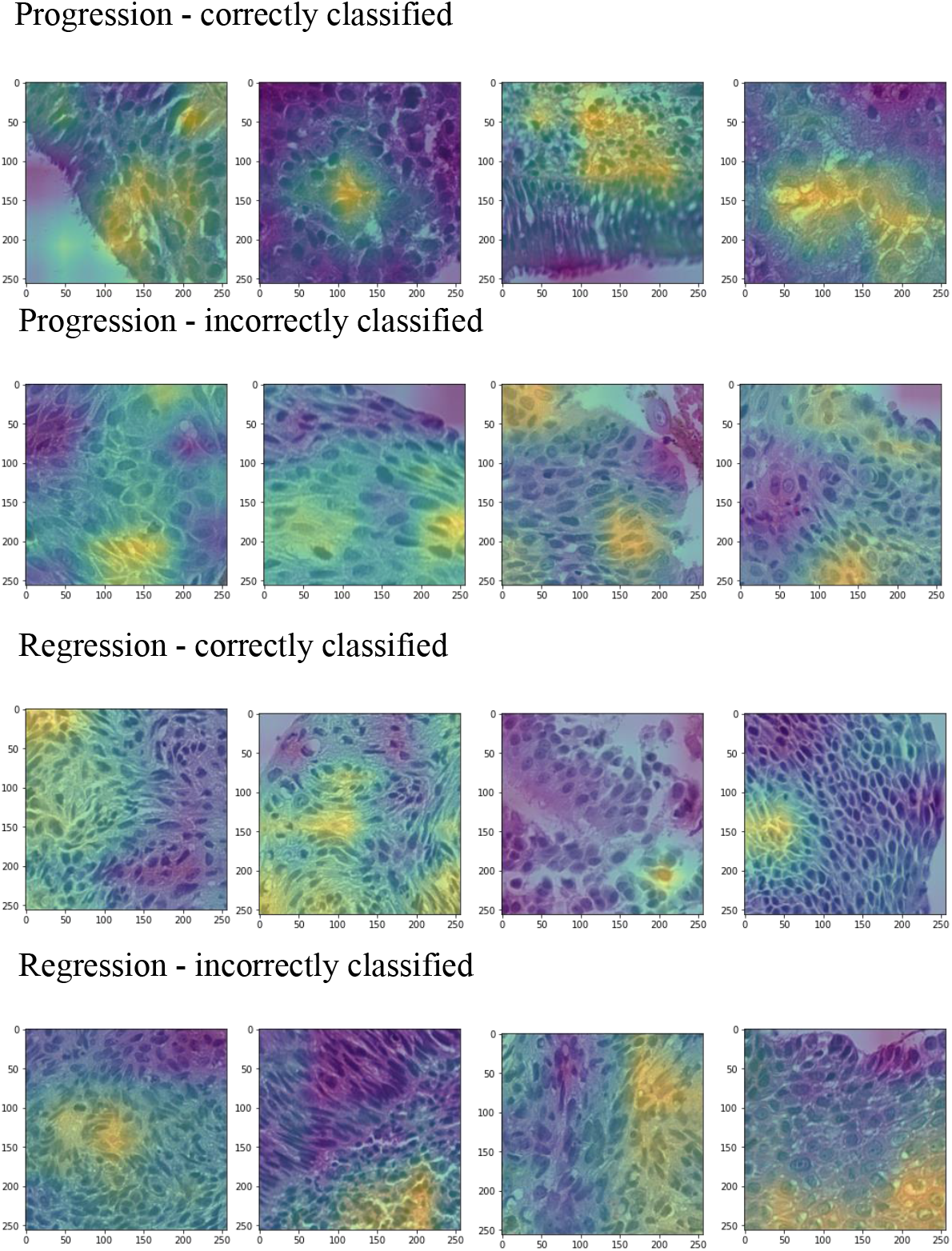
Class activation maps for progression and regression

## Discussion

The inability to predict the evolution of SCIS creates problems in the management of patients that are diagnosed with these lesions. Considering the fact that 30% of SCIS regress and never become invasive carcinomas, overtreating these lesions is an issue. Currently, there are no histologic criteria in place for making such a prediction. Even though previous studies have managed to find molecular markers that are predictive for progression, molecular assays are still difficult to implement at scale, especially in the developing world, where the incidence of SCC of the lung is higher (21). The ability to upload photos of SCIS lesions taken from slides to a cloud-based prediction algorithm could facilitate access in resource-poor settings. Here, we present a DL based approach which relies only on H&E stained WSI that managed to reach a per-tile classification AUC of 0.78 and sensitivity (recall) of 0.94. Our approach proved to be very capable of detecting the progression of SCIS to SCC, as shown by the high sensitivity and the clusters suggested by the t-SNE analysis. The F1 score of 0.84 highlights the fact that the model is accurate even when factoring in the imbalanced dataset. Given that our model was trained using a very small dataset by the standards of a DCNN, we believe that DL is a promising tool for predicting the progression of preinvasive lesions.

We also generated class activation maps in order to understand the inner workings of the model. Explainability is a key aspect to take into account when it comes to the adoption of artificial intelligence (AI) in pathology. In the future, the creation of feedback loops between the pathologist and the models used will be vital for increasing accuracy, decision-making transparency and providing a pathologist-centered AI model. Various methods of visualizing the way DL models generate their predictions could also aid the pathologist in the discovering new morphologic clues.

The study has certain limitations. Our dataset only consisted of 112 WSI out of which we managed to extract 11130 images. DL models are very data hungry and their performance increases when provided with more data (22). To create even more powerful predictive models, more data would be needed. However, sizeable cohorts like the one that our data came from are very rare as they require performing invasive procedures as well as a long follow-up period. Collaboration and pooling of smaller in-house cohorts would be needed to generate larger datasets on which even more accurate models could be trained.

## Conclusion

In conclusion, our study suggests that DL could be used to predict the evolution of SCIS. We believe this approach could be used to generate similar predictions about other preinvasive precursor lesions in other organs, using only H&E stained WSI. As more and more countries adopt screening programs and larger databases of these lesions will become available, DL has the potential to become part of the screening toolkit and to be a high throughput and low-cost way of providing patients and clinicians with more prognostic information.

## Acknowledgements

We would like to thank Pennycuick et al. for making their dataset publicly available (14).

## References

1. Sung H, Ferlay J, Siegel RL, Laversanne M, Soerjomataram I, Jemal A, et al. Global Cancer Statistics 2020: GLOBOCAN Estimates of Incidence and Mortality Worldwide for 36 Cancers in 185 Countries. CA A Cancer J Clin. 2021 May;71(3):209–49.

2. Barta JA, Powell CA, Wisnivesky JP. Global Epidemiology of Lung Cancer. Annals of Global Health. 2019 Jan 22;85(1):8.

3. Sánchez-Danés A, Blanpain C. Deciphering the cells of origin of squamous cell carcinomas. Nat Rev Cancer. 2018 Sep;18(9):549–61.

4. Herbst RS, Morgensztern D, Boshoff C. The biology and management of non-small cell lung cancer. Nature. 2018 Jan;553(7689):446–54.

5. Thakrar RM, Pennycuick A, Borg E, Janes SM. Preinvasive disease of the airway. Cancer Treatment Reviews. 2017 Jul;58:77–90.

6. Pipinikas CP, Kiropoulos TS, Teixeira VH, Brown JM, Varanou A, Falzon M, et al. Cell migration leads to spatially distinct but clonally related airway cancer precursors. Thorax. 2014 Jun;69(6):548–57.

7. The National Lung Screening Trial Research Team. Reduced Lung-Cancer Mortality with Low-Dose Computed Tomographic Screening. N Engl J Med. 2011 Aug 4;365(5):395–409.

8. Teixeira VH, Pipinikas CP, Pennycuick A, Lee-Six H, Chandrasekharan D, Beane J, et al. Deciphering the genomic, epigenomic, and transcriptomic landscapes of pre-invasive lung cancer lesions. Nat Med. 2019 Mar;25(3):517–25.

9. Abiodun OI, Kiru MU, Jantan A, Omolara AE, Dada KV, Umar AM, et al. Comprehensive Review of Artificial Neural Network Applications to Pattern Recognition. IEEE Access. 2019;7:158820–46.

10. Echle A, Rindtorff NT, Brinker TJ, Luedde T, Pearson AT, Kather JN. Deep learning in cancer pathology: a new generation of clinical biomarkers. Br J Cancer. 2021 Feb 16;124(4):686–96.

11. Iizuka O, Kanavati F, Kato K, Rambeau M, Arihiro K, Tsuneki M. Deep Learning Models for Histopathological Classification of Gastric and Colonic Epithelial Tumours. Sci Rep. 2020 Dec;10(1):1504.

12. Coudray N, Ocampo PS, Sakellaropoulos T, Narula N, Snuderl M, Fenyö D, et al. Classification and mutation prediction from non-small cell lung cancer histopathology images using deep learning. Nat Med. 2018 Oct;24(10):1559–67.

13. Wulczyn E, Steiner DF, Xu Z, Sadhwani A, Wang H, Flament-Auvigne I, et al. Deep learning-based survival prediction for multiple cancer types using histopathology images. Hsieh JCH, editor. PLoS ONE. 2020 Jun 17;15(6):e0233678.

14. Pennycuick A, Teixeira VH, AbdulJabbar K, Raza SEA, Lund T, Akarca AU, et al. Immune Surveillance in Clinical Regression of Preinvasive Squamous Cell Lung Cancer. Cancer Discovery. 2020 Oct 1;10(10):1489–99.

15. Williams E, Moore J, Li SW, Rustici G, Tarkowska A, Chessel A, et al. Image Data Resource: a bioimage data integration and publication platform. Nat Methods. 2017 Aug 1;14(8):775–81.

16. Jeremy George P, Banerjee AK, Read CA, O’Sullivan C, Falzon M, Pezzella F, et al. Surveillance for the detection of early lung cancer in patients with bronchial dysplasia. Thorax. 2007 Jan 1;62(1):43–50.

17. He K, Zhang X, Ren S, Sun J. Deep Residual Learning for Image Recognition. In: 2016 IEEE Conference on Computer Vision and Pattern Recognition (CVPR) [Internet]. Las Vegas, NV, USA: IEEE; 2016 [cited 2022 Oct 25]. p. 770–8. Available from: http://ieeexplore.ieee.org/document/7780459/

18. Paszke A, Gross S, Massa F, Lerer A, Bradbury J, Chanan G, et al. PyTorch: An Imperative Style, High-Performance Deep Learning Library. 2019 [cited 2022 Oct 25]; Available from: https://arxiv.org/abs/1912.01703

19. Krizhevsky A, Sutskever I, Hinton GE. ImageNet classification with deep convolutional neural networks. Commun ACM. 2017 May 24;60(6):84–90.

20. Zhou B, Khosla A, Lapedriza A, Oliva A, Torralba A. Learning Deep Features for Discriminative Localization. 2015 [cited 2022 Oct 25]; Available from: https://arxiv.org/abs/1512.04150

21. Cheng TYD, Cramb SM, Baade PD, Youlden DR, Nwogu C, Reid ME. The International Epidemiology of Lung Cancer: Latest Trends, Disparities, and Tumor Characteristics. Journal of Thoracic Oncology. 2016 Oct;11(10):1653–71.

22. Alom MZ, Taha TM, Yakopcic C, Westberg S, Sidike P, Nasrin MS, et al. A State-of-the-Art Survey on Deep Learning Theory and Architectures. Electronics. 2019 Mar 5;8(3):292.

